# Thermal acclimation and resultant developmental adaptation offsets environmental temperature effects on tail muscle mechanics in larval zebrafish

**DOI:** 10.64898/2026.08.14.744876

**Authors:** Andrew F. Mead, Marcus A. Zimmermann, Michael J. Previs, David M. Warshaw

## Abstract

Environmental temperature strongly influences muscle contractile mechanics and locomotor performance in ectotherms, yet animals routinely develop across a range of temperatures while maintaining effective movement. We tested the hypothesis that developmental temperature induces compensatory changes in the intrinsic mechanical properties of the muscles that power the fast-start escape response in larval zebrafish (*Danio rerio*). Larvae were reared at 25°C, 28°C, or 32°C, and contractile properties of intact tail myotomal muscles were measured across experimental temperatures. Acute changes in experimental temperature strongly affected twitch kinetics, particularly relaxation rate (Q_10_ = 2.1), resulting in substantial changes in twitch duration. In contrast, rearing temperature produced adaptive changes that opposed these acute thermal effects. At a common experimental temperature, muscles from cold-reared larvae exhibited faster intrinsic relaxation and greater force production during shortening at a physiologically relevant velocity, whereas warm-reared larvae showed slower relaxation and reduced shortening force. As a result, twitch kinetics were largely normalized when measurements were made at each group’s rearing temperature, reducing the apparent thermal sensitivity of relaxation rate (Q_10_ = 1.1). To identify molecular correlates of these functional adaptations, we performed label-free quantitative LCMS proteomic analysis. Cold rearing increased the abundance of Sarco/Endoplasmic Reticulum Calcium-ATPase (SERCA) proteins, driven primarily by elevated *atp2a1* expression, while warm rearing reduced the abundance of the major parvalbumin isoforms *pvalb1* and *pvalb2*. These changes implicate remodeling of intracellular calcium handling as a mechanism underlying thermal compensation of muscle function. Together, our results demonstrate that developmental temperature modifies the intrinsic mechanical properties of larval zebrafish muscle in ways that counteract the direct effects of environmental temperature, thereby preserving the timing and power-generating capacity required for fast-start escape performance.

## Introduction

Vertebrate skeletal muscle is an adaptable biological machine that relies on highly conserved cellular components to power a wide diversity of animal movements. Much of our understanding of this versatility comes from comparative physiological studies, which have revealed how muscle mechanical output relates to whole-body biomechanics across variations in body size, mode of locomotion, and environmental temperature (Rome *et al*., 1988; Johnston, 1991; James and Johnston, 1998; Wakeling and Johnston, 1998; Mendoza, Moen and Holt, 2023). In ectothermic animals such as fish, environmental temperature directly influences the biochemical reaction rates that underlie muscle contraction, strongly affecting locomotor performance (Bennett, 1984; Rall and Woledge, 1990; James, 2013). As a result, swimming efficiency and performance typically decline outside an optimal temperature range, driven in part by a mismatch between the intrinsic mechanical properties of muscle and the biomechanical demands of locomotion (Rome, 1995; Johnston, 2006; Oufiero and Whitlow, 2016).

Many fish species can partially compensate for the impact of acute temperature changes on muscle function through thermal acclimation over both short and long timescales, and during development (Johnston, 2006; Scott and Johnston, 2012). A substantial component of thermal acclimation arises from adjustments to the intrinsic mechanical properties of muscle cells themselves by varying the expression of muscle gene paralogs with distinct biochemical properties (Johnston and Temple, 2002; Watabe, 2002). However, the cellular and molecular mechanisms that enable such compensation, particularly in muscles operating at extreme contractile speeds, remain incompletely understood (Mendoza, Moen and Holt, 2023).

The emergence of zebrafish (*Danio rerio*) as a genetic and physiological model organism, together with a growing body of biomechanical and genomic data, offers new opportunities to dissect the molecular mechanisms of muscle mechanical performance and temperature adaptation. This opportunity is particularly compelling during early larval stages, when zebrafish exhibit one of the fastest locomotor behaviors observed in vertebrates: the “fast-start” escape response. By approximately 5 days post-fertilization, larvae perform a maneuver consisting of an initial unilateral body bend (the “C-start”) followed by a burst of rapid alternating tail beats (Domenici and Blake, 1997). In zebrafish larvae, this behavior displaces the animal by several body lengths within tens of milliseconds, with individual tail-beat cycles lasting as little as ∼12 msec (Voesenek, Muijres and van Leeuwen, 2018). Such rapid movements require myotomal tail muscles capable of precise temporal control of activation and relaxation, as well as the ability to generate force at extremely high shortening velocities to power effective hydrodynamic thrust (Voesenek, Muijres and van Leeuwen, 2018; Voesenek *et al*., 2019). *Ex vivo* mechanical measurements of intact larval zebrafish tails have revealed some of the fastest contractile properties reported for vertebrate muscle (Mead *et al*., 2020, 2024), and further indicate that these muscles operate near their functional limits during fast-start behavior (Mead *et al*., 2020; Ravel *et al*., 2025).

Zebrafish are typically reared and studied at 28°C, a temperature at which larval fast-start performance and muscle contractile properties are well characterized (Voesenek *et al*., 2019; Mead *et al*., 2020). However, they can be reared successfully in the laboratory across a 25–32°C temperature range, and likely experience broader thermal variation in natural environments (Scott and Johnston, 2012). In this study, we test the hypothesis that rearing zebrafish at temperatures below and above 28°C (i.e., 25°C and 32°C, respectively) induces compensatory changes in muscle mechanical performance, specifically (i) the kinetics of *in situ* larval tail twitch force and (ii) the force–velocity relationship so as to preserve the optimal relationship between muscle contractile properties and fast-start biomechanics.

Our results show that for larvae reared at 28°C, twitch kinetics and force-velocity performance are strongly influenced by the temperature at which these physiological parameters were measured. Amazingly, rearing larvae at cooler (25°C) or warmer (32°C) temperatures leads to developmental compensation so that these same physiological parameters when measured at the rearing temperature become equivalent to that of larvae reared at 28°C. Using quantitative proteomics, we find significant changes in the abundance and isoform composition of intracellular calcium handling proteins (i.e., the sarcoplasmic reticular calcium ATPase pump, SERCA, and the soluble calcium binding protein, parvalbumin) which may contribute to the compensatory adaptation in the larval tail muscle mechanical performance. Such developmental compensation may maintain the tail muscle’s ability to generate the rapid fast-start necessary to escape predators.

## Results and Discussion

### Developmental plasticity minimizes the thermal sensitivity of twitch duration

In many vertebrate locomotory muscle systems, the duration of active force production during a contraction is primarily controlled by the duration of a train of membrane action potentials triggered by motor neurons. In contrast, EMG recordings and *in situ* muscle mechanics indicate that during the zebrafish larval fast-start escape response each tail beat results from a single evoked action potential (Buckingham and Ali, 2004; Mead *et al*., 2020). In this type of contraction, the duration of force generation is determined by the speed of processes intrinsic to the muscle cells, rather than the duration of the neural stimulus. These processes include biochemical rates associated with excitation-contraction (E-C) coupling, force development, and relaxation, which are known to be sensitive to temperature, with temperature coeficients (Q_10_, defined as the factor by which a rate changes in response to a temperature increase of 10°C) spanning 1 to >5 (Rall and Woledge, 1990; James, 2013). Thus, acute changes in environmental temperature within the normal physiological range would at a minimum be expected to affect the duration of the muscle’s response to a neural input during a fast-start. If so, continuous exposure to higher or lower temperatures within that range during development may result in a counteracting adaptive response to maintain a constant duration of force generation.

To observe how acute and rearing temperatures affect how larval myotomal muscles respond to a unitary stimulus, we used a modified classical approach to measure the transient rise in isometric force resulting from a sub millisecond electrical stimulation, known as a ‘twitch’ (see Methods as originally detailed in Mead *et al*., 2020) (Fig. 1). Briefly, tails were mounted between a servo motor and force transducer in a temperature-controlled bath. After a short period of equilibration, preparations were stimulated with a single 0.4 msec field stimulus pulse, and the resulting twitch was analyzed for peak force, twitch duration (t_50-50_; time from 50% of peak force during force development to 50% peak force during relaxation), rate of force development (+dF/dT t_20-80_; 20–80% peak force development), and relaxation rate (-dF/dT t_80-20_; 80–20% peak force) (Fig. 1A).

**Figure 1.**
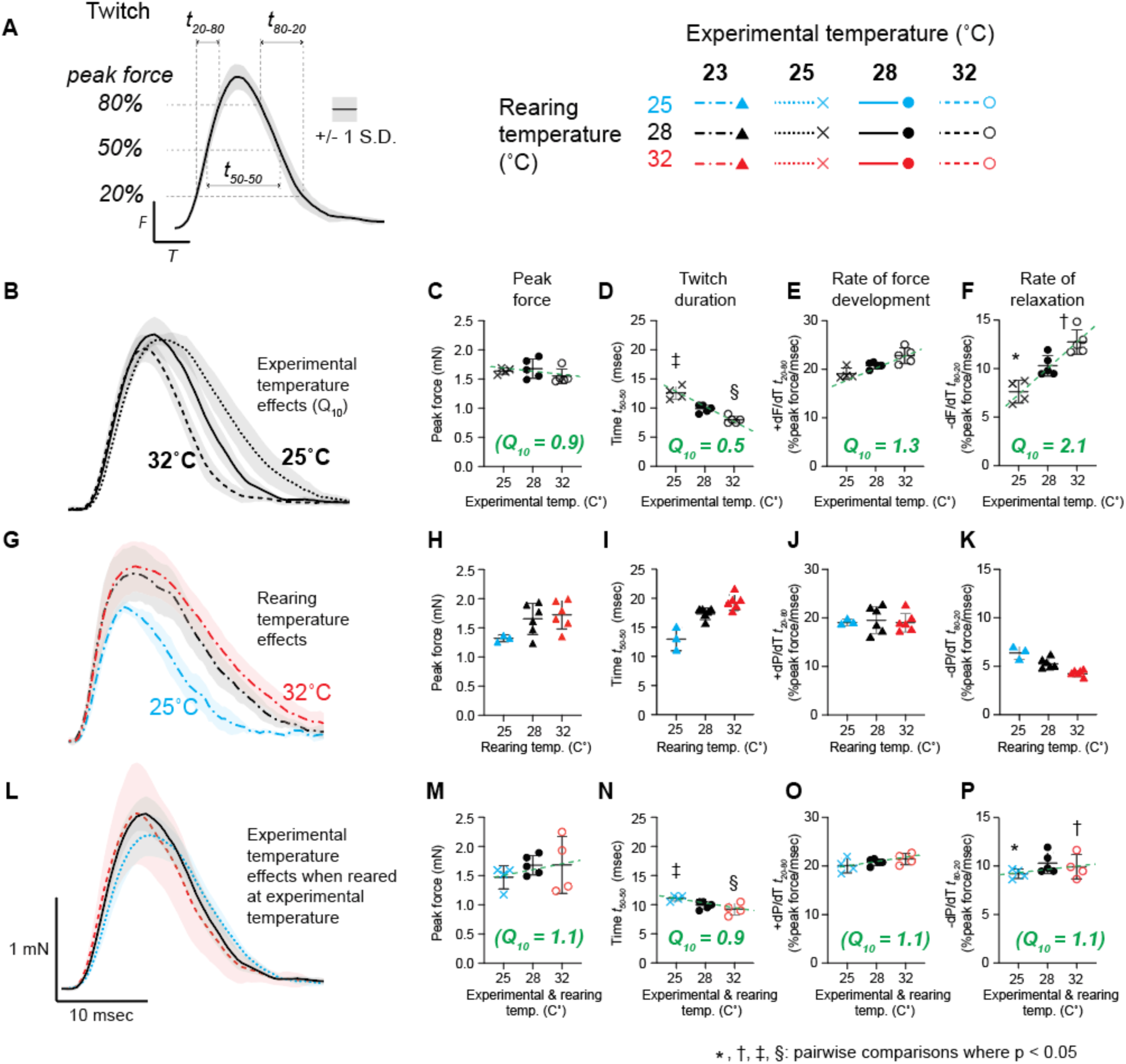
Effects of experimental and rearing temperature on twitch kinetics in intact 5 dpf zebrafish tails. **(A)** Legend showing measured parameters of the ‘twitch’ force transient. Traces are averaged from twitches evoked by 0.4 msec field stimulation pulses (n = 3-6 animals). Shaded regions depict +/- 1SD. Line and symbol ***color*** depict rearing temperature for panels B-P. Line and symbol ***type*** depict experimental temperature for panels B-P. **(B-F)** Effects of experimental temperature on twitch properties of tails from larvae reared at 28°C. Experimental temperature had significant effects (1-way anova) on twitch duration (D, p = 0.002), rate of force development (E, p = 0.0003), and relaxation (F, p = 0.0002), but not peak force (C, p = 0.3). Temperature coefficient estimates (Q_10_) are calculated as described in Materials and Methods. Parentheses are used to denote Q_10_ values derived from properties with non-significant effects by 1-way anova (i. e. peak force, C). **(G-K)** Effect of rearing temperature on twitch properties measured at 23°C. Rearing temperature was positively correlated with twitch duration (I, p = 0.0001) and negatively correlated with relaxation rate (K, p = 0.0001), but had no effect on peak force (H, p = 0.083) or rate of force development (J, p = 0.9). **(L-P)** Rearing at experimental temperature prior to experimentation neutralized the effect of experimental temperature alone. Tails reared at 25°C had significantly shorter twitch durations (‡; p = 0.033), and faster relaxation rates (*, p = 0.032) when measured at 25°C than those reared at 28°C. Those reared at 32°C had longer durations (§, p = 0.046) and slower relaxation rates (†, p = 0.006) at 32°C than those reared at 28°C.

To measure the acute effects of temperature, twitches were collected from larvae reared at 28°C with the experimental chamber set to 25°, 28°, or 32°C (Fig. 1 B-F). Peak twitch force was insensitive to experimental temperature over the studied range (1-way ANOVA, p = 0.3; Fig. 1 B,C), although variable temperature effects on peak twitch force have been reported depending on muscle type and species (Bennett, 1984). This variability has been attributed to the counteracting effects of temperature on the rate of force-generating actomyosin cross bridge formation versus the duration of the activating cytosolic [Ca^2+^] transient (Bennett, 1984; Rome, 1995). However, twitch duration was negatively correlated with experimental temperature, equivalent to a Q_10_ of 0.5 (p = 0.002; Fig. 1 B, D). This resulted from the temperature sensitivities of the rates of force development (p = 0.003, Q_10_ = 1.3; Fig. 1 B, E), and, especially, relaxation (p = 0.0002; Q_10_ = 2.1; Fig. 1 B, F). These results broadly agree with values previously reported from a range of species and muscle types, with the exception being the rate of force development, which typically shows a larger temperature dependence than reported here (Rall and Woledge, 1990; James, 2013).

To observe adaptive effects on the twitch following prolonged exposure to environmental temperature, larvae were reared to 5 dpf at 25°, 28°, or 32°C and twitch parameters measured at a common experimental temperature of 23°C (Fig. 1 G-K), which was chosen to facilitate isovelocity force measurements (see below). Again, peak twitch force did not differ significantly among rearing groups (p = 0.083, Fig. 1 G, H). However, contrary to the effects of acute temperature, rearing temperature had a significant positive effect on twitch duration (p = 0.0001, Fig. 1 G, I). This was driven primarily by a negative correlation between rearing temperature and relaxation rate (p = 0.0001; Fig. 1 G, K), as rearing temperature had no effect on the rate of force development (p = 0.9; Fig. 1 G, J). Interestingly, this effect of rearing temperature on relaxation rate (equivalent to Q_10_ = 0.5, Fig. 1 G, K) was in the equal and opposite direction to that when larvae were reared at 28°C and the experimental temperature varied between 25° and 32°C (Q_10_ = 2.1, Fig. 1 B, F).

These data suggest that changes in rearing temperature as the larvae develop alter muscle physiology in an attempt to counteract the acute thermal sensitivity of muscle function so that the fast-start escape response is maintained during periods of prolonged changes in environmental temperature. If so, then measuring twitch responses at experimental temperatures matched to each group’s rearing temperature (25°, 28° or 32°C; Fig. 1 L-P) should result in equivalent twitches. This in fact was the case as all measured twitch parameters for the larvae reared at 25_°_C and then measured at 25°C were no different than those reared at either 28°C or 32°C and measured at 28°C and 32°C, respectively. By normalizing the twitch duration across environmental temperatures, this form of developmental plasticity may preserve the ability of larval muscle to cycle at the frequencies biomechanically optimal for fast-starts.

### Rearing temperature inversely affects the force-velocity relationship

While larval muscle twitch force generated under isometric conditions is not sensitive to temperature, locomotory power during the fast-start is a mechanical property known to be strongly temperature dependent and depends on the force generated while muscle fibers are shortening (Voesenek, Muijres and van Leeuwen, 2018). The relationship between force and shortening velocity is described by the force–velocity curve (Fig. 2 A), in which force declines as shortening velocity increases, from maximal force under isometric conditions (F_max_) to minimal or no force at maximal shortening velocity (V_max_). Muscle power, being the product of force and shortening velocity, is maximal at intermediate shortening velocities (Fig. 2 A). During swimming, fish muscle operates within a narrow range of velocities where power and efficiency are optimal (V/V_max_ between 0.18 and 0.35 (Rome *et al*., 1988; Rome, 1995). Since both V_max_ and maximal power (P_max_) have reported Q_10_ values of 1.5 or greater (Rome, 1995; James, 2013), changes in environmental temperature have the potential to create a mismatch between optimal muscle performance and whole-body biomechanics during the fast-start. We therefore hypothesized that, as is the case with twitch kinetics that larval myotomal muscle undergoes compensatory changes in its force-velocity properties.

**Figure 2.**
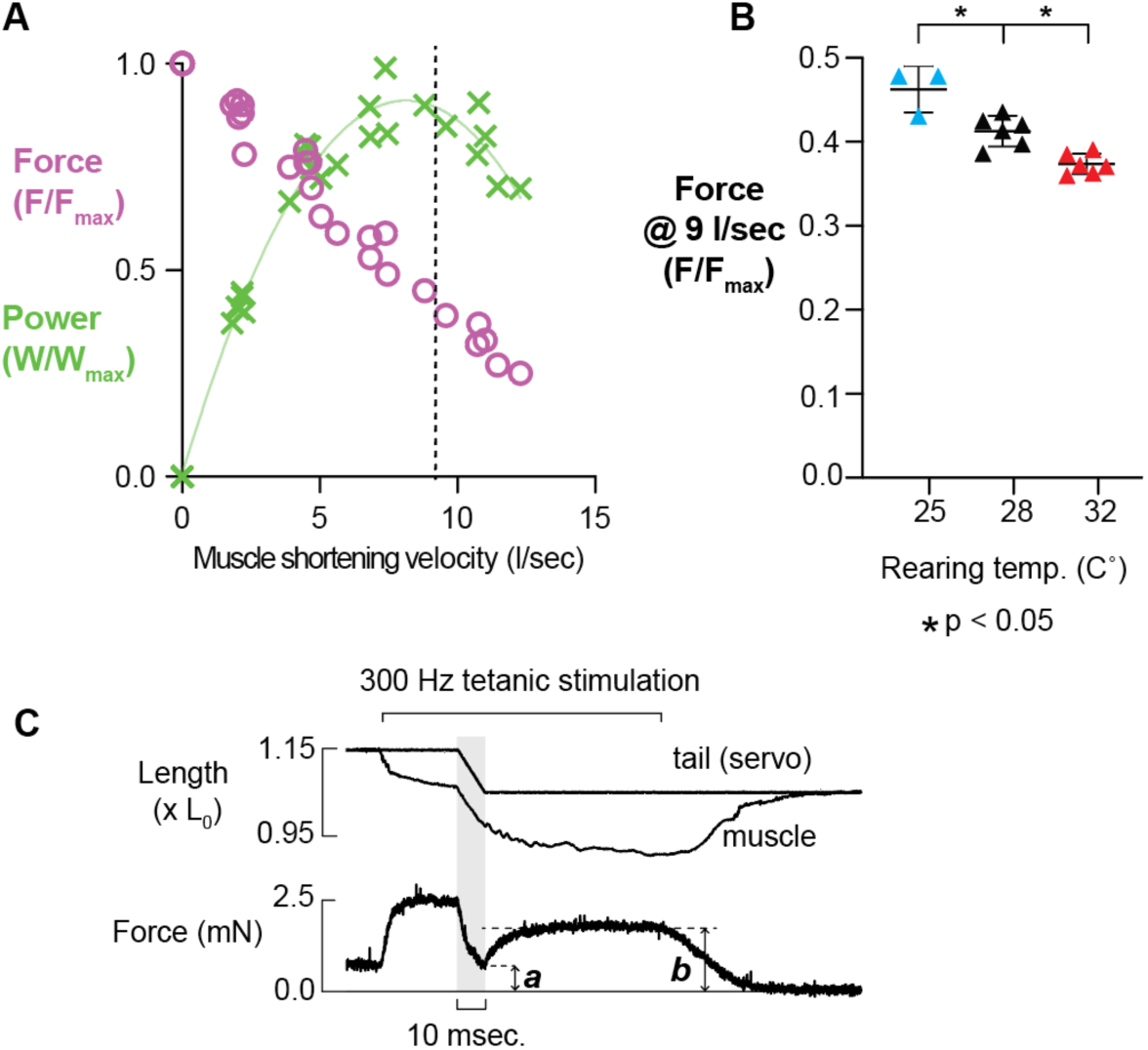
Effect of rearing temperature on myotomal muscle force during shortening. **(A)** Force-velocity data (purple data points) and power-velocity (green data points) from 5 dpf larval tails at 23°C, reproduced from Mead 2024. Dashed vertical line denotes velocity (i.e. 9 l/sec.) at which data in panel ‘B’ was collected. **(B)** The effect of rearing temperature on force during shortening at 9 l/sec, the velocity at which P_max_ is produced. 25°C-reared tails developed more force during shortening (p = 0.013), and 32°C-reared tails less (p = 0.0014), than 28°C-reared tails. **(C)** Representative data traces from an isovelocity maneuver used to generate data in panel ‘B’. Tails were tetanized using a 300 Hz train of 0.4 msec field stimulation pulses and allowed to shorten at 9 l/sec for 10 msec by means of the servo. Muscle length and velocity were measured by tracking myoseptal position using a high-speed camera as described in Mead 2020 and 2024. F/F_max_ values in panels ‘A’ and ‘B’ are calculated as the ratio of minimum shortening force (*a*) to recovered isometric force (*b*).

Due to the extreme speed of larval muscle and the limited temporal response of our recording instruments, we are unable to record the entire force-velocity relationship. Therefore, we previously reported partial force–velocity relationships of 5 dpf zebrafish larvae reared at 28°C but then studied at 23°C to slow the muscle responses and thus accommodate our instrumentation using *in situ* isovelocity shortening maneuvers (Fig. 2 A, purple data points) (Mead *et al*., 2024). These data indicate that P_max_ is generated near 9 muscle lengths/sec at 23°C (Fig. 2 A, green data points). To investigate whether rearing temperature leads to compensating effects on force while shortening at a velocity near P_max_, we reared larvae to 5 dpf at 25°, 28°, or 32°C and conducted isovelocity tests at 9 lengths/sec at a common experimental temperature of 23°C (Fig. 2 B, C). Briefly, tails were mounted as described above and stimulated with a 300 Hz train of pulses to achieve steady state tetanic contraction (Fig. 2 C). During tetanus, myotomal muscles were allowed to shorten at 9 lengths/sec for 10 msec by means of the length servo. The force recorded at the end of the isovelocity maneuver was compared to the subsequent isometric force after recovery in order to control for the dependence of muscle force on muscle length (Mead *et al*., 2020).

At this velocity, larvae reared at 28°C generated an F/F_max_ of 0.41 +/- 0.02, consistent with previous measurements (Mead *et al*., 2024). Tails from 25°C reared larvae generated significantly greater relative force while shortening at 9 lengths/sec (F/F_max_ of 0.46 +/- 0.03; p = 0.013) than tails from 28°C reared larvae. The opposite was true of tails from 32°C reared larvae (F/F_max_ of 0.37 +/- 0.01; p = 0.0014) (Fig. 2 B). As with relaxation kinetics (Fig. 1 G-K), the direction of the adaptive response of force during an isovelocity maneuver may serve to offset the acute temperature effects expected for V_max_ and P_max_ (Rome, 1995; James, 2013), suggesting that developmental temperature induces compensatory shifts in the force-velocity properties to maintain appropriate power for the fast-start response over a range of developmental temperatures.

### Label-free quantitative liquid chromatography mass spectrometry (LCMS): rearing temperature affects abundance and isoform composition of key Ca^2+^-handling myotomal muscle proteins

To determine whether the adaptive effects of rearing temperature on the observed contractile properties are accompanied by molecular remodeling, we used label-free quantitative LCMS optimized for zebrafish larval muscle (O’Leary et al., 2019; Wood et al., 2022; Mead et al., 2024) to examine the abundance and isoform composition of muscle proteins with well-characterized roles in excitation and contraction. Briefly, tails from larvae reared at 25, 28, or 32°C were digested with trypsin and analyzed by label-free LCMS (Materials and Methods; Mead *et al*., 2024). As in our previous study (Mead *et al*., 2024), our analysis identified peptides that mapped to products from individual genes (unique peptides) as well as peptides that mapped to multiple annotated proteins (shared peptides) encoded by related genes with regions of conserved sequence (File S1). Unique peptides were used to quantify individual protein isoforms, whereas shared peptides provided a measure of the combined abundance of closely related proteins. To quantify the abundance of individual protein isoforms, the average abundance of the top two or three unique peptides mapping to each protein was normalized to the average abundance of the top three peptides shared among all identified sarcomeric myosin heavy chain isoforms (encoded by: *myhz1.1, myhz1.3, myhz2, and smyhc1*). This was done to provide a measure of each protein’s abundance relative to the amount of myotomal muscle analyzed and to minimize the influence of variation in sample recovery and LCMS signal among samples. The same quantification was performed for closely related proteins with multiple shared peptides. We then compared normalized abundance values for individual and related proteins from the 25°C and 32°C groups to those of 28°C controls. A complete list of individual proteins and paralogous groups for which at least two peptides were identified, together with abundance values and statistics, is included in Table S1.

The extent and duration of twitch force generation is highly dependent on the timecourse of the cytosolic [Ca^2+^] transient following a stimulus (Berchtold, Brinkmeier and Müntener, 2000; Rome, 2006; Mead *et al*., 2017). With activation, Ca^2+^ released from the sarcoplasmic reticulum (SR) binds to the troponin-tropomyosin complex, exposing myosin binding sites on actin, allowing myosin to attach and generate force. In the present study, rearing temperature primarily affected twitch duration through changes in relaxation rate. Therefore we hypothesized that this effect would be accompanied by changes in proteins responsible for reducing cytosolic [Ca^2+^] to resting levels after activation, specifically the sarco/endoplasmic reticulum calcium ATPases (SERCA) pumps and the soluable calcium binding parvalbumins (Fig. 3).

**Figure 3.**
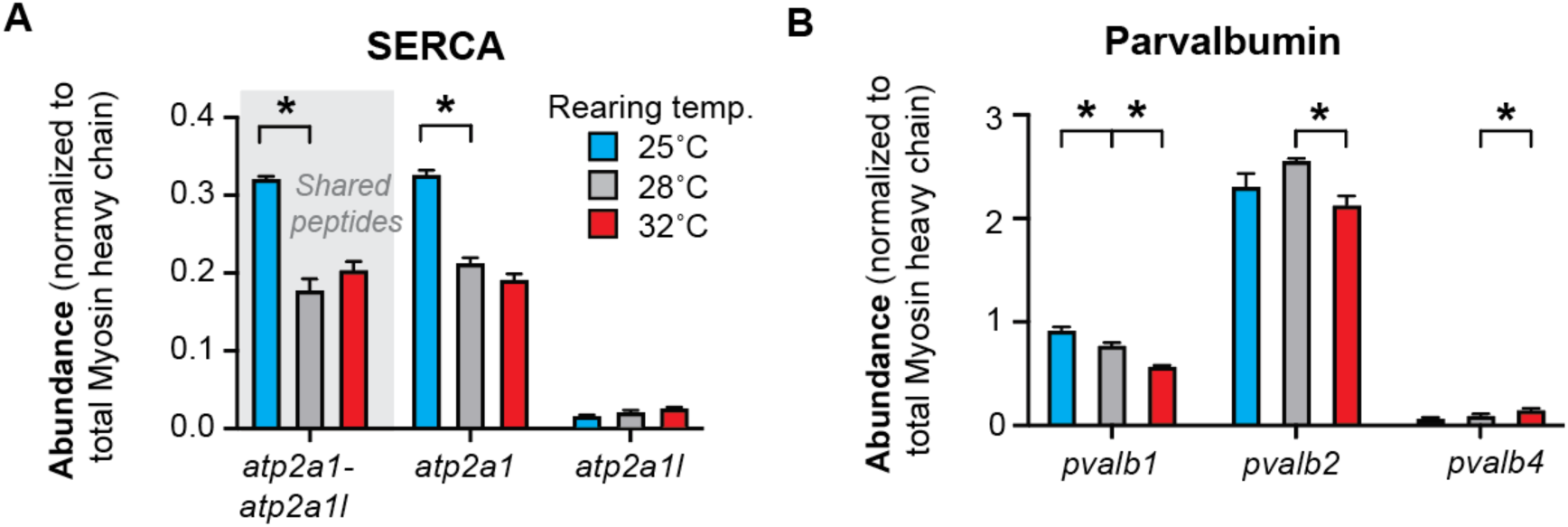
Effects of rearing temperature on abundance and isoform composition of SERCA and parvalbumin. **(A, B)** Abundances of peptdies associated with SERCA and parvalbumin gene products as a function of rearing temperature. Abundances are normalized to total myosin heavy chain abundance. The number of peptides identified for each gene product are shown in parentheses. In the case of SERCA, peptides were identified with sequences shared between individual isoforms. Note: direct comparison of the abundance of individual isoforms or between individual isoform and shared peptides are not possible due to the different ionization efficiencies of individual peptides. n = 3 individual 5 dpf larval tails per temperature group. Asterisks denote differences where p < 0.05.

SERCA, located in the sarcoplasmic reticular (SR) membrane, pumps Ca^2+^ ions from the cytosol into the SR lumen. Therefore, muscle relaxation rate depends on the overall abundance of SERCA as well as the various isoforms having different Ca^2+^ affinities (Berchtold, Brinkmeier and Müntener, 2000; Periasamy and Kalyanasundaram, 2007; Mead *et al*., 2017). Here, we identified peptides from two SERCA isoforms present in all tail samples, encoded by *atp2a1* and *atp2a1l* genes, which are orthologs of mammalian fast-twitch SERCA1 (Fig. 3 A, Table S2). Rearing at 25°C versus 28°C resulted in a nearly 2-fold increase in the molar abundance of peptides shared by both isoforms relative to myosin heavy chain (p < 0.0001) (Fig. 3 A, Table S2). Analysis of isoform-specific peptides indicates that the increase in shared peptides is primarily driven by increases in *atp2a1* expression (p < 0.0001), whereas the less-abundant *atp2a1l* expression was unchaged (p = 0.058) (Fig. 3 A, Table S2). These results suggest a substantially increased capacity for Ca²⁺ resequestration into the SR in cold-acclimated muscle, consistent with our twitch measurements after cold-rearing, which show an increased intrinsic relaxation rate (Fig. 1 B, F). Interestingly, rearing at 32°C did not produce the opposite effect; total SERCA abundance, as indicated by shared peptides, was unchanged (Fig. 3 A, Table S2; p = 0.071), as were isoform specific peptides associated with *atp2a1* (p = 0.065), and *a2p2a1l* (p=0.065).

In muscles with fast relaxation rates, especially those of fish and amphibians, the soluble calcium binding protein, parvalbumin, also impacts the [Ca^2+^] transient time course (Arif, 2009). Levels of parvalbumin expression is broadly correlated with contractile speed in fishes (Rome, 2006; Schoenman *et al*., 2010; Campion *et al*., 2012) and increased expression is associated with cold acclimation in smelt (Woytanowski and Coughlin, 2013). As with SERCA, parvalbumin isoforms differ in their Ca²⁺ affinity, suggesting the impact of parvalbumin on cytosolic free [Ca^2+^] is isoform specific (Arif, 2009; Schwaller, 2020). We identified unique parvalbumin isoform peptides encoded by three genes (*pvalb1, pvalb2, pvalb4*) (Fig. 3 B, Table S2). Larvae reared at 25°C compared to rearing at 28°C showed an increase in the molar abundance of *pvalb1* peptides relative to myosin heavy chain (p = 0.003), and no change in *pvalb2* (p = 0.058), or *pvalb4* peptide abundance (p = 0.061). For 32°C reared larvae compared to those reared at 28°C both *pvalb1* (p = 0.001) and *pvalb2* (p = 0.001) decreased while *pbalv4* increased (p = 0.011). Unfortunately, lack of sufficient shared peptides prevented a direct comparison of total parvalbumin abundance between rearing groups. However, our data do show that *pvalb1* and *pvalb2* were among the most abundant proteins of all those identified in our samples, underscoring the importance that even small shifts in their relative abundance may have on Ca^2+^ handling, especially if these isoforms are functionally distinct (Table S1).

Together, these results indicate that the rearing-temperature effects on twitch duration, and particularly twitch relaxation rate, are accompanied by coordinated changes in the abundance and isoform composition of proteins that determine the duration of the activating Ca²⁺ transient. Notably, these molecular responses were strongly asymmetric. Cold rearing produced substantial increases in the abundance of proteins that promote Ca²⁺ sequestration, whereas warm rearing resulted primarily in shifts in parvalbumin isoform composition with no significant change in total SERCA abundance. Thus, although both developmental temperatures produced compensatory changes in muscle function, they appear to do so through quantitatively different patterns of molecular remodeling rather than equal and opposite adjustments. Such asymmetric thermal responses have been observed in other ectotherm systems and may reflect the greater physiological challenge imposed by maintaining rapid contractile kinetics at low temperatures (Johnston and Temple, 2002; Rome, 2006).

### Limitations of the study

Although our LCMS characterization identified rearing temperature shifts in abundance and isoform composition of critical Ca^2+^-handling proteins that may account for the compensatory changes in the twitch relaxation rate, our LCMS data unfortunately do not provide definitive insight to the rearing temperature effect on proteins that impact the force-velocity parameters. Specifically, steady state mechanical properties of force, velocity, and power can be related to the various myosin heavy chain isoforms and their accompanying essential and regulatory light chains (Bottinelli and Reggiani, 2000). For example, larval zebrafish fast myotomal muscle cells have been shown to express six closely related fast-type myosin heavy chain genes (*myhz1.1*, *myhz1.2*, *myhz1.3, myhz2, myhc4, and myha*) in a distribution that varies along the length of the tail (Nord *et al*., 2014). Of these, we identified peptides mapping to *myhz1.1, myhz1.3,* and *myhz2* in all samples. However, only 10 out of 99 of these peptides were unique to a single isoform, the remainder being shared by two or more isoforms (Table S1, File S1) likely due to the high degree (≥96%) of sequence similarity between fast-type isoforms (Nord *et al*., 2014). Thus, we were unable to resolve whether rearing temperature affected the relative abundance of individual fast-type myosin heavy chain isoforms. While not thought to contribute to the fast-start response, slow muscle fibers constitute about 7% of myotomal muscle cross-sectional area at 5 dpf (Mead *et al*., 2020). Noteably, the abundance of peptides unique to the slow type myosin heavy chain *smyhc1* was not affected by rearing temperature (Table S1), indicating that functional changes were not due to shifts in the relative amount of fast vs. slow type muscle. In addition, key myosin binding partners, such as titin and myosin binding protein-C and -H have been shown to impact myosin molecular function (Fusi *et al*., 2016; Li *et al*., 2019; Mead *et al*., 2024). However, we did not detect changes in their abundance or isoform composition (Table S1) even though post-translational modifications, such as phosphorylation, of these myosin binding partners may occur in response to rearing temperature and thus potentially contribute to contractile plasticity (Krysiak *et al*., 2018; Li *et al*., 2019; Robinett *et al*., 2019).

Although, the significant changes in contractile parameters with rearing temperature were interpreted as adaptations to the thermal environment, temperature alone is known to accelerate larval development (Kimmel *et al*., 1995). Therefore, larvae reared at 32°C could be considered developmentally more advanced (i.e., ‘older’) at 5 dpf than those reared at 28°C, with those reared at 25°C being ‘younger’. Despite this, we saw no difference between rearing temperature groups in larval body length (excluding tail fin) (p = 0.18) or body height measured at the point of the anal vent (p = 0.11) at 5 dpf (Fig, S1 A-C). To control for the contractile differences between rearing groups being due to ‘biological age’ rather than temperature acclimation directly, we compared twitch and force-velocity responses at 25°C experimental temperature from larvae reared at 28°C to 4 dpf (n = 3) and 6 dpf (n = 3) (Fig. S1 D-H). If the functional effects of rearing temperature that we observed (Figs. 1 G, 2 B) were caused due to biological age, we would expect to see similar changes with chronological age, which we did not. In fact, no significant effect of chronological age was observed between groups in twitch force (p = 0.58), twitch duration (p = 0.78), rate of force development (p = 0.61), rate of relaxation (p = 0.43), or force during shortening (p = 0.16). Therefore, temperature-dependent aging effects cannot explain the changes in larval tail muscle mechanics observed with rearing temperature.

Additional mechanisms, beyond those examined here, are also relevant to fast-start performance and its sensitivity to environmental temperature. *In vivo*, myotomal muscles operate under dynamic strains that include rapid shortening and re-lengthening imposed by activation of contralateral tail muscle fibers, conditions which can alter the timecourse of force development and relaxation (Josephson, 1993; Mendoza, Moen and Holt, 2023). These dynamic properties may help explain why twitch durations measured here (∼10 msec) under isometric conditions exceed the shortening phase of an 80-Hz tail-beat cycle (∼6 msec) (Voesenek, Muijres and van Leeuwen, 2018; Mead *et al*., 2020). In addition, muscle fiber-extrinsic factors such as fiber orientation, body and fin morphology, and neural activation patterns could contribute to whole-animal compensation for temperature-dependent changes in muscle mechanics (Voesenek, Muijres and van Leeuwen, 2018).

**Figure S1.**
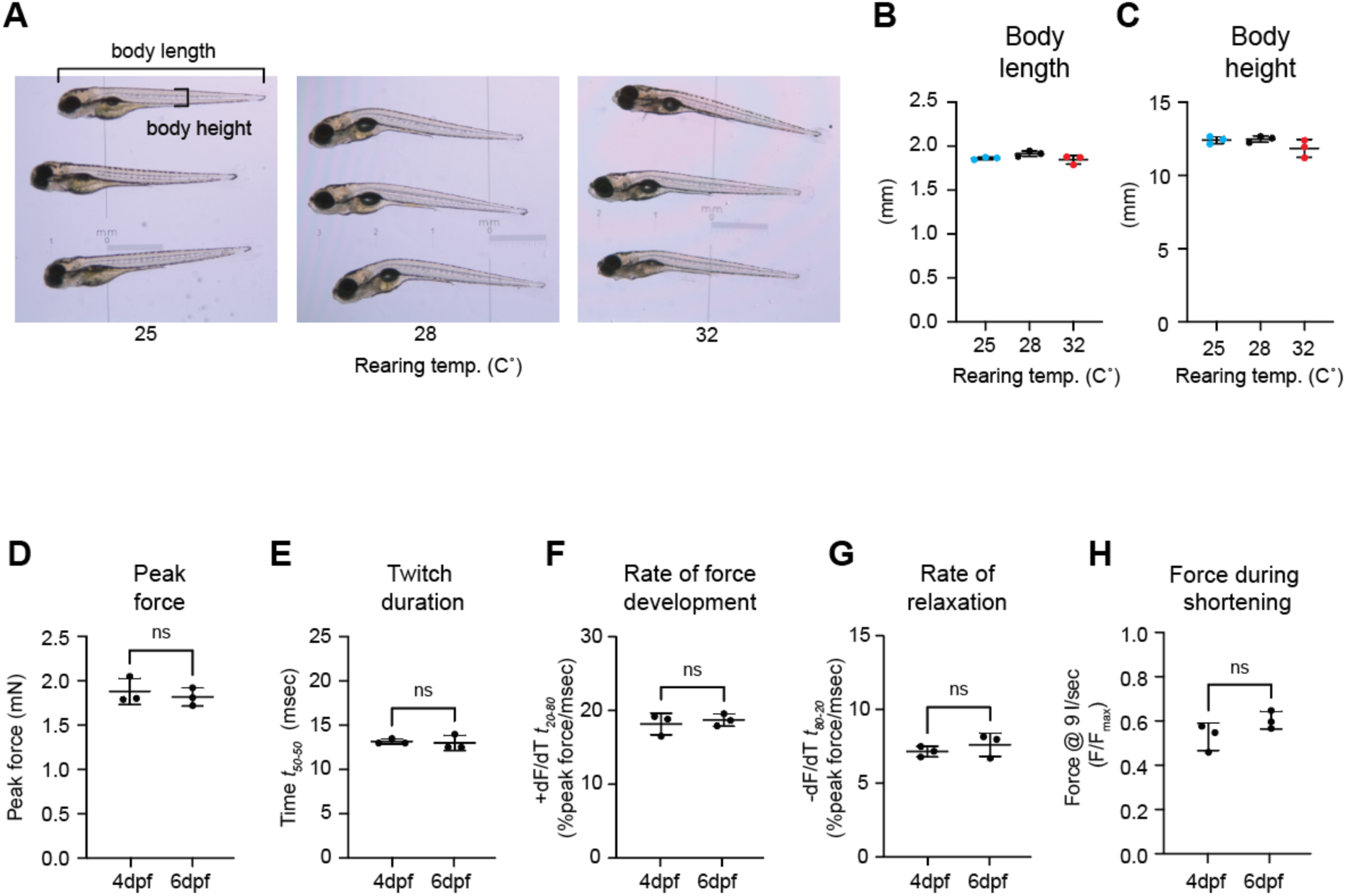
**A-C**: Comparison of body size between 5 dpf larvae reared at 25°, 28°, and 32°C. No difference was seen in body length (A, B) (n = 3, p = 0.18) or body height (A, C) (n = 3, p = 0.11). **D-H**: Comparison of *in vitro* mechanical parameters of tail muscles from larvae reared at 28°C to 4dpf and 6dpf and measured at 25°C. No significant difference was seen in peak twitch force (D, n = 3, p = 0.58), twitch duration (E, n = 3, p = 0.78), rate of force development (F, n = 3, p = 0.61), rate of relaxation (G, n = 3, p = 0.43), or force during shortening (H, n = 3, p = 0.16).

## Conclusion

Ectothermic animals must cope with the direct effects of environmental temperature on muscle function and locomotor performance. Although thermal adaptation has been widely documented in fish, its mechanistic basis in fast-contracting locomotor muscles remains poorly understood. Using larval zebrafish as a model system, we show that rearing temperature during early development induces compensatory changes in intrinsic muscle mechanical properties that are critical for fast-start escape response performance. Specifically, rearing at cooler temperatures produced muscles with faster intrinsic relaxation kinetics and enhanced force generation at physiologically relevant shortening velocities, whereas rearing at warmer temperatures resulted in slower relaxation and reduced force generation at physiologically relevant shortening velocities. Notably, these developmental adaptations are opposite to the acute thermal effects on muscle contractility, effectively minimizing the impact of environmental temperature on the fast-start escape response that is so critical for larval survival. Quantitative proteomic analysis identified coordinated changes in the abundance and isoform composition of key excitation–contraction coupling proteins, including SERCA and parvalbumin, which likely contribute to the observed modulation of relaxation kinetics. Together, these findings highlight the zebrafish model as a powerful system for uncovering conserved mechanisms of contractile regulation and thermal adaptation in vertebrate muscle.

## Materials and Methods

### Animals and Husbandry

All experiments were approved by the Institutional Animal Care and Use Committee at the University of Vermont, and were in compliance with the Guide for the Use and Care of Laboratory Animals published by the National Institutes of Health. Larvae used for functional and proteomic studies came from AB strain breeding pairs acquired from ZIRC (RRID:ZIRC_ZL1). Embryos and larvae were maintained in E3 medium (5 mM NaCl, 0.17 mM KCl, 0.33 mM CaCl_2_, 0.33 mM MgSO_4_, 0.0001% methylene blue (Sigma-Aldrich)). Sibling embryos were divided at random into three groups and placed in icubators set to 25, 28, or 32°C within an hour of fertilization at a density of 50 embryos per dish. For all studies, larvae were maintained at their rearing temperature until immediately before euthanasia and either mounting in the mechanics rig or preparation for LCMS.

### Muscle Mechanics

Isometric twitch and isovelocity force measurements were performed using previously described methods (Mead *et al*., 2020, 2024), with modifications specific to the present study described below. Larvae were selected at random from each rearing group, euthanized by tricaine overdose (0.05% in E3), and transferred to Ringer’s solution (117.2 mM NaCl, 4.7 mM KCl, 1.2 mM KH_2_PO4, 1.2 mM MgCl_2_, 2.5 mM CaCl_2_, 25.2 mM NaHCO_3_, 11.1 mM glucose; oxygenated and equilibrated to pH 7.4 with a mixture of 95% O_2_ and 5% CO_2_). Tails were removed immediately caudal to the swim bladder and mounted between a force transducer (400C; Aurora Scientific Aurora, Ontario, Canada) and linear servo motor (MC1; SI, Heidelberg, Germany) using spring clamps spaced 1 mm apart. The anal vent was positioned equidistant from the attachment points. Preparations were imaged at 5,000 frames/s using a CCD camera (IL5; Fastec Imaging, Knoxville, TN) with custom optics (0.95 µm/pixel). Force, motor position, and camera timing were recorded at 40 kHz using custom data-acquisition software written in IGOR (WaveMetrics, Portland, OR).

Following mounting, preparations were returned to their in vivo length (L_0_) to remove strain introduced during attachment and then stretched to 1.05 × L_0_. Preparation temperature was maintained at 23, 25, 28, or 32°C by continuously perfusing oxygenated Ringer’s solution (0.7 ml/min) through the chamber using a custom countercurrent heat exchanger. Chamber temperature was verified at the beginning and end of each experiment. Preparations were allowed to equilibrate for 20 min before stimulation. Twitches were elicited with a single 0.4-ms, 7-V pulse using a MyoPacer stimulator (IonOptix, Westwood, MA), with the attachment clamps serving as electrodes.

For isovelocity measurements, preparations were stretched to 1.15 × L0 at 0.5 preparation lengths/s and tetanized with a 100-ms train of 0.4-ms pulses at 300 Hz. Forty milliseconds after the onset of tetanus, preparations were shortened from 1.15 to 1.05 × L0 at constant velocity. The minimum force reached during shortening was expressed as a proportion of maximal active isometric force, measured 20 ms after the shortening ramp when force had returned to a plateau. Three sequential isovelocity contractions were performed for each preparation and averaged.

Q_10_ estimates were calculated using mean twitch parameter values at 25° and 32°C according to the equation:

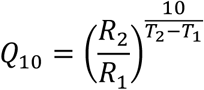

where *R* is the value of a temperature-dependent rate and *T* is the experimental temperature at which that rate was measured.

### Larval Imaging

Live imaging of larvae (Fig. S1) was peformed at 5 dpf using a Nikon SMZ800 dissecting stereomicroscope (Nikon instruments, Melville, NY) with Insight 2 camera and Spot Basic software (Diagnostic Instruments, Sterling Heights, MI) Fish were oriented over a calibration slide. Body measurements were made manually in Fiji software (Image J) (Schindelin *et al*., 2012).

### Protein Quantification

Quantitative liquid chromatography mass spectrometry (LCMS) was performed using a previously described label-free workflow (O’Leary *et al*., 2019; Mead *et al*., 2024), with the modifications and analysis specific to the present study described below. Individual 5 dpf tails were placed in 1.5-ml microcentrifuge tubes containing 150 µl 0.1% RapiGest SF Surfactant (Waters Corporation). Samples were heated at 50°C for 45–60 min, reduced with 0.75 µl 1 M dithiothreitol at 100°C for 10 min, and alkylated with 22.5 µl 100 mM iodoacetamide in 50 mM ammonium bicarbonate for 30 min in the dark at 22°C. Proteins were digested overnight at 37°C with 5 µg trypsin in 50 mM ammonium bicarbonate. RapiGest was subsequently cleaved with formic acid and trifluoroacetic acid as previously described (Mead et al., 2024), and samples were dried, reconstituted in 0.1% trifluoroacetic acid, and centrifuged at 18,800 × g for 5 min before transfer of the supernatant to mass spectrometry (MS) vials.

Peptides were separated by UHPLC using an XSelect UPLC HSS T3 column (3.5 µm, 1.0 × 150 mm; Waters Corporation) and analyzed on a Q Exactive Hybrid Quadrupole-Orbitrap mass spectrometer (Thermo Fisher Scientific) using data-dependent MS/MS acquisition. The five most abundant ions were selected for fragmentation. Peptide spectra were searched against the Danio rerio proteome database (UP000000437; UniProt, downloaded February 2015) using SEQUEST in Proteome Discoverer 2.2 (RRID: SCR_014477). Variable modifications included N-terminal methionine loss, N-terminal acetylation, carbamidomethylation of cysteine, oxidation of methionine and proline, and phosphorylation of serine, threonine, and tyrosine. The Minora Feature Detector was used to identify chromatographic peaks corresponding to peptide spectral matches across samples.

Label-free quantification followed O’Leary *et al*., (2019). Peak areas reported by Proteome Discoverer were exported for analysis in RStudio (RRID: SCR_000432). For each protein or group of proteins with shared peptides, the two or three most abundant peptides were selected based on abundance across experimental groups (File S1, Tab: ‘Top 1-3 peptides-all hits’). Proteins and groups of proteins with only one peptide, or that were not identified in all samples, were eliminated from subsequent analysis. Isoform-specific and group-specific abundances were calculated from the averaged abundance of these peptides (File S1, Tab: ‘Averaged Top 2 or 3 Peptides’) and normalized to the averaged abundance of the three most abundant peptides shared among all identified sarcomeric myosin heavy chain isoforms (File S1, Tab: Normalization; Table S1). Normalized values for larvae reared at 25°C and 32°C were then compared with those from the 28°C group (Tables S1, S2; Fig. 3).

### Statistcal Analysis

Data are reported as means +/- 1 standard deviation. Statistical analysis was performed using Prism 11 (Graphpad Software, Boston, MA). Effects of experimental temperature and rearing temperature on twitch parameters (Fig. 1) were analyzed by one-way analysis of variance (ANOVA). Pairwise comparisons of twitch durations and relaxation rates from samples reared at 25°C vs. 28°C and measured at 25°C, and samples reared at 32°C vs. 28°C and measured at 32°C (Fig. 1 F, D, N, P) were peformed using one-tailed, unpaired t-tests. Pairwise comparisons of force during shortening from samples reared at 25°C or 32°C vs. samples reared at 28°C were analyzed using two-tailed, unpaired t-tests (Fig. 2). Age effects on functional parametrs (Fig. S1 C-G) were analyzed using two-tailed, unpaired t-tests. Effects of rearing temperature on larval body size (Fig. S1 A,B) were analyzed by one-way analysis of variance (ANOVA). For exploratory proteomic analysis (Fig. 3; Table S1), planned pairwise comparisons (25°C vs. 28°C and 32°C vs. 28°C) for each identified protein and shared peptide group were performed using multiple unpaired two-tailed t-tests with false discovery rate controlled using the two-stage step-up method (Benjamini, Krieger and Yekutieli, 2006) (Table S1). Proteins and peptide groups selected a priori (Table S2) were analyzed using multiple t-tests followed by Holm correction for multiple testing. Statistical significance was defined as p < 0.05.

